# Potential for Climate Change induced extinction of the Sky Island Species Mount Graham Red Squirrel (*Tamiasciurus hudsonicus grahamensis*)

**DOI:** 10.64898/2026.05.13.725054

**Authors:** Eric Gibson, Michael Kantar, Bryan Runck

## Abstract

Sky islands are high-elevation ecosystems surrounded by lowland habitats that create isolated environments with distinct climatic conditions. These factors have driven the evolution of many endemic species, separated from their larger, contiguous populations. An Individual-Based Model (IBM) was used to simulate population dynamics by modeling the behaviors and interactions of *Tamiasciurus hudsonicus* grahamensis (Mount Graham Red Squirrel) a subspecies of the American red squirrel (*Tamiasciurus hudsonicus*) that is endemic to the Pinaleño Mountains in southeastern Arizona. This approach can help predict future population trends based on historical species data leading to better conservation decisions. Using species-specific ecological preferences—including temperature, precipitation, and vegetation indices (NDVI)—an IBM was developed to simulate population dynamics and spatial distribution projections through 2100. Climate change projections, based on the best- and worst-case scenarios outlined in the 2014 National Climate Assessment, were incorporated to assess potential future population trends under changing environmental conditions. The population faces a 45-62% probability of extinction by 2100, with a significant risk of extinction within the next 50 years. A translocation experiment was conducted to evaluate the viability of relocating individuals to the Chiricahua Mountains, another sky island with a larger habitable area. However, the risk of extinction remains even higher (87-89%) due to environmental disturbances affecting both the Chiricahua and Pinaleño regions. This highlights the challenges of conservation efforts in the face of climate change and emphasizes the need for targeted management strategies to preserve this critically endangered subspecies.

## Introduction

Climate change is profoundly altering biogeography and sky islands offer a focused scope to examine the impacts of this changes (Love, 2023). Island biogeography provides a theoretical framework to understand the impact of human societal problems on other species. Specifically, as habitat gets more fragmented due to climate change more species are impacted. The effect will be particularly acute for already endangered species. Sky Islands are natural phenomena that have provided geographic isolation and refuge from historic climate fluctuations (Love et al., 2023). Typically, sky islands emerge in mountainous regions, with species moving up and down the slope during times of climate stress and having periods of species mixing during times of ambient climate (Love et al., 2023). The makeup of any specific sky island will influence the characteristics of the species that live there. Understanding the environmental parameters of any sky island as well as the species that live there allows for accurate predictions about both historic and potential future phenotypic changes. For example, many species which have moved to islands have gained or lost body mass when separated from their ancestors due to changes in resource availability and competition (Lomolino 1985). Sky islands offer essential data while minimizing monetary and temporal costs, making them indispensable tools for advancing environmental science and biology.

This study focuses on the Mount Graham red squirrel (*Tamiasciurus hudsonicus grahamensis*), a critically endangered subspecies of the American red squirrel (*Tamiasciurus hudsonicus*) confined to the Pinaleño Mountains in southeastern Arizona. This mountain range is part of the Madrean Sky Islands, isolated mountain ecosystems surrounded by desert lowlands. The squirrel’s habitat is primarily high-elevation mixed-conifer and spruce-fir forests, which provide food resources, shelter, and suitable microclimatic conditions. This decrease in habitat has occurred through forest fires, changes in insect pest distributions, and increased competition with other species. *T. h. grahamensis* prefers high-elevation montane forest habitat, so sky islands with larger areas of montane forests will support a healthier population of the species. Additionally, as climate change negatively affects the size of montane forest habitat due to increased desertification and wildfire risk, environmental support will likely decline and less migration and an overall decrease in population size will occur (Wood 2010).

Understanding changing habitat and demographics can help with on the ground decision support for conservation practitioners (McLane et al., 2011; Guisan, et al., 2013). Recently there have been increased efforts to make species distribution models more accessible and relevant to conservation managers in particular future potential ranges (Frans et al., 2022). These distributions which include temperature, precipitation and ground cover, provide necessary inputs for understanding population changes using an agent-based model (Tang et al., 2010; Watkins et al., 2015). Therefore, the goal of this study was to predict the future distribution of sky islands in south-west North America and to predict how these habitat changes will impact the *T. h. grahamensis* population. This information will provide an opportunity to understand what interventions can be made to prevent the extinction of this vulnerable species. The Mount Graham Red Squirrel provides critical insights into how mammal species in the Pinaleño Mountains and across montane forest habitats, may respond to climate change. By analyzing shifts in the population size and distribution of *T. h. grahamensis*, predictions can extend to broader semi-arid regions worldwide.

## Material and Methods

### Data Acquisition

Sky island locations were sourced from the United States Geological Survey’s (USGS) Earth Explorer platform, which provides a wide range of raster datasets derived from both satellite-based and in-situ collection methods. Temperature and precipitation data for 30-year climate normals were obtained from the WorldClim database at 1 km resolution (Fick and Hijmans, 2017). Normalized Difference Vegetation Index (NDVI) was retrieved from the Earth Resources Observation and Science (EROS) Center’s Visible Infrared Imaging Radiometer Suite (eVIIRS). This satellite-derived raster dataset has a spatial resolution of 375 meters Monthly NDVI data for 2023 was averaged over 7- or 14-day periods, depending on weather conditions. While higher spatial resolution NDVI rasters were considered, cloud cover— particularly during the winter months—compromised data accuracy in the study area. Consequently, a lower resolution dataset was adopted to ensure data functionality and reliability within the model. Species occurrence data for *T. h. grahamensis* were obtained from the Global Biodiversity Information Facility (GBIF.org (19 November 2024) GBIF Occurrence Download https://doi.org/10.15468/dl.szwdhb. The full workflow can be found in **Figure 1**.

**Figure 1.**
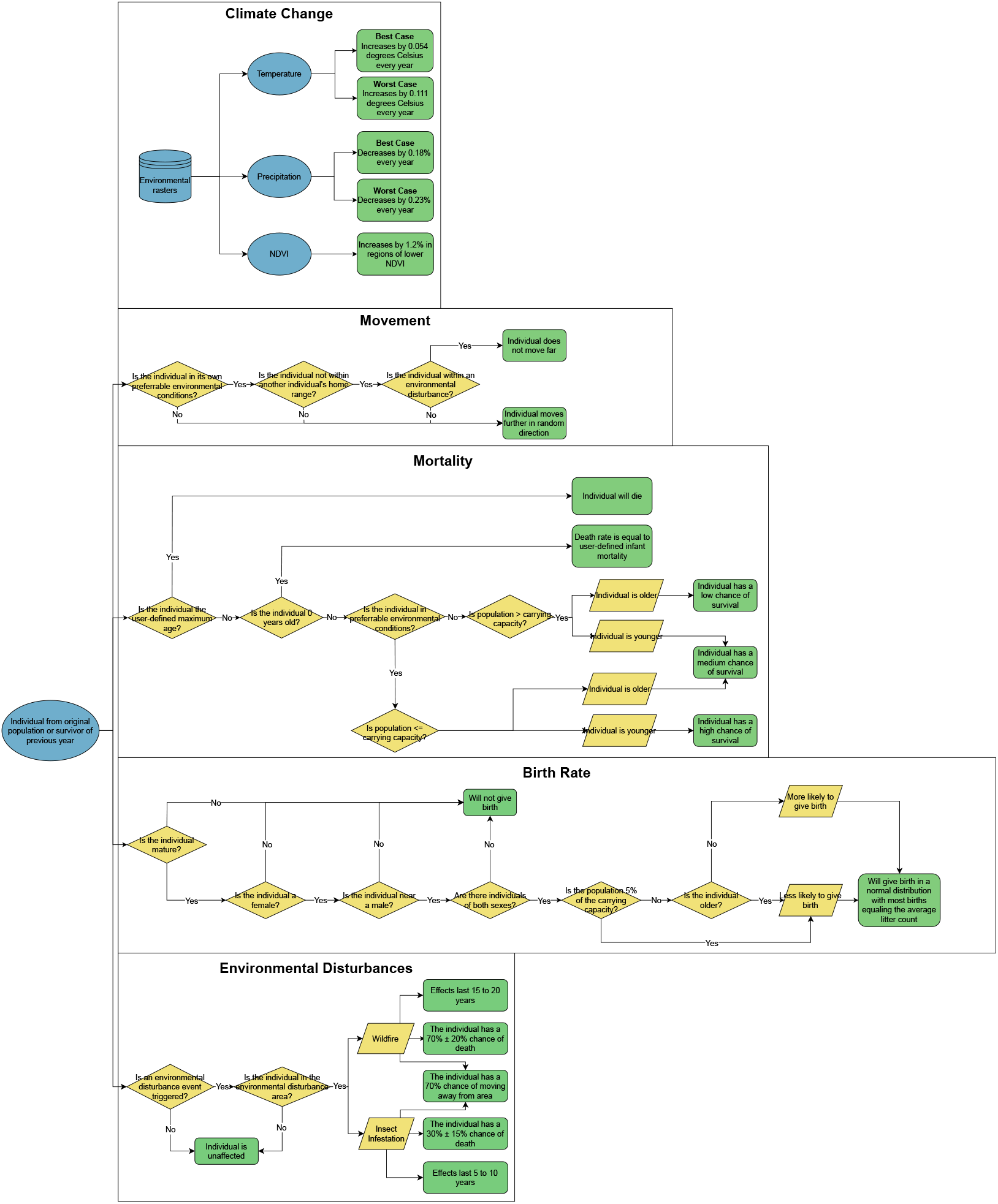
Data flow diagram illustrating the structure of the Individual-Based Model. The model incorporates climate change effects, individual movement, birth rates, mortality rates, and environmental disturbances. Every individual is then tracked for location, age, and sex, which inform their unique life-history characteristics and annual distributions. Environmental disturbances in the region primarily include wildfires and insect infestations. Wildfires typically affect areas ranging from 1,000 to 50,000 acres and occur approximately every 10 years, as indicated by recent fire history in the Pinaleño Mountains (Koprowski et al., 2006; Koprowski et al., 2008; Rushton et al., 2018; U.S. Fish and Wildlife Service, 2024). Insect infestations, which generally occur every 30 years, tend to impact nearly the entire coniferous forest—areas characterized by the highest NDVI values within the model.

### Sky Island Delineation

To determine sky island boundaries and the environmental conditions influencing species distribution, monthly temperature, precipitation, and NDVI raster data were utilized. After the initial calculation each raster dataset was reclassified according to sky island thresholds, a maximum average monthly temperature of 12.5°C, a minimum yearly precipitation of 500 mm and a minimum monthly NDVI of 0.3, with cells meeting these criteria classified as part of a sky island. Given that sky islands lack clear, delineated boundaries, an accuracy assessment was conducted to validate the model (Grother et al., 2005). The assessment utilized occurrence points of *T. h. grahamensis*. First, occurrence points in the southwestern United States and northern Mexico were identified within areas meeting the sky island thresholds and an equivalent number of random points were generated across the same study area. The counts of squirrel occurrence points within the modeled sky islands were compared to the counts of random points using a t-test. The *T. h. grahamensis* inhabits sky islands characterized by higher precipitation, higher vegetation productivity, and lower temperatures compared to the surrounding areas. Adverse environmental conditions outside these favorable habitats can significantly increase mortality due to resource scarcity, such as a lack of food, water, or shelter in trees (Menzies et al., 2020). Their survival is constrained by a specific temperature range, as regions in the southern United States are generally too warm for *T. h. grahamensis* to persist outside the sky islands (Johannesdottir, 2017).

### Individual-Based Model

An IBM was developed to simulate population dynamics and spatial distribution of the *T. h. grahamensis*. The model incorporates the effects of climate change, habitat suitability, and demographic parameters on individual movement, reproduction, and mortality (**Table 1**). It also evaluates the potential migration, emigration, and population trends under future climate scenarios. To initialize the population, 144 individuals were selected based on the most recent 2023 data provided by the University of Arizona (Koprowski, 2024). The spatial distribution of the initial population was modeled using a combination of known *T. h. grahamensis* occurrence data and randomly generated points. The distribution was defined by taking the mean geographic location of all recorded *T. h. grahamensis* occurrences were calculated and created a buffer zone of approximately 23.4 kilometers (equivalent to 0.25 decimal degrees). Points representing individuals were randomly placed within the buffer zone, constrained to regions meeting specific environmental criteria. Suitable areas were defined as those with an average annual NDVI > 3500 and an average annual precipitation of > 60 millimeters. These criteria were used to identify habitat regions where individuals were likely to find essential resources, providing a realistic and ecologically informed starting distribution for the simulation.

**Table 1.** Definitions of life-history characteristics, population distributions, and environmental conditions influencing the annual birth and death rates of T. h. grahamensis. The “Standard” column indicates whether the statistic is supported by findings from other scientific studies. *See equation variables in Supplemental Materials.

| <u>*T. h. grahamensis* Simulation Statistics</u> |  |  |  |
| --- | --- | --- | --- |
| Trait | Statistic/Equation | Standard | Reference |
| Age of maturity | 1 year | Y | Descamps et al., 2008 |
| Average litter size | ~4 offspring | N | Hayssen, 2008 |
| Litter size range | 1 to 8 offspring | Y | Hayssen, 2008 |
| Percentage of females who give birth per year | ~85% | N | Koprowski, 2005 |
| Infant Mortality | 75% | Y | Halvorson et al., 1983 |
| Average lifespan | 5 years | Y | Steele, 1998 |
| Maximum age | 10 years | Y | Steele, 1998 |
| Age-based birth rate* | $C_b = (0.85 \times \frac{1}{1 + (0.5 \times (a - a_m))^e})$ | N | Descamps et al., 2008 |
| Age-based mortality rate* | $C_d = \frac{1}{1 + (-0.9 \times (a - a_l))^e}$ | N | No reference |
| Population-based mortality rate* | $w_p = e^{exp_p} \times P^3$ | N | U.S. Fish and Wildlife Service, 2024 |
| Wildfire mortality rate | 70% ± 20% | N | U.S. Fish and Wildlife Service, 2024 |
| Insect infestation mortality rate | 30% ± 15% | N | U.S. Fish and Wildlife Service, 2024 |

#### Initial Age Distribution

The age distribution of the *T. h. grahamensis* has not been formally documented. However, based on expert insights (U.S. Fish and Wildlife Service, 2024), it is hypothesized that the population comprises a greater proportion of younger individuals aged between 2 and 4 years compared to other age groups. This assumption provides a foundational basis for initializing age distribution in the simulation. Additionally, stochasticity was implemented to initialize the ages within the population slightly differently with each simulation.

#### Individual Preferences

Individual environmental preferences were determined using monthly climatic raster data and occurrence records for the *T. hudsonicus* across Arizona and New Mexico. Due to the limited occurrence data and small sample size of the *T. h. grahamensis*, preference distributions were derived from regional *T. hudsonicus* occurrence data, including records for the Mount Graham subspecies. This approach was chosen because the *T. hudsonicus* is the closest relative to the *T. h. grahamensis* and shares similar ecological requirements. To establish preference distributions, raster cell values corresponding to *T. hudsonicus* occurrence points were extracted for each environmental variable—precipitation, temperature, and NDVI. These extracted values formed the basis of the initial distribution of individual preferences. A Kernel Density (KDE) Model was created for each environmental variable, capturing the range and probability of preference values observed across the species’ occurrence data. Every new individual, whether part of the initial population or born during the simulation, was assigned a random preference for monthly temperature, precipitation, and NDVI. Preferences were drawn probabilistically from the distributions generated by the KDE Models, ensuring that individual preferences reflected the environmental conditions favored by the species. This approach allowed the model to dynamically simulate individual variation in environmental preferences while accounting for the ecological niche of *T. h. grahamensis* within the broader context of the *T. hudsonicus* population.

#### Individual Movement

The movement of individuals within the model is determined based on individual environmental preferences. The maximum movement per year was set to approximately 679.8 meters per individual, as reported for *T. h. grahamensis* by Koprowski et al. (University of Arizona). This distance translates to approximately 0.00726 decimal degrees horizontally and 0.00611 decimal degrees vertically. To calculate the yearly movement, the model first computes the difference between the environmental values of the current location and the individual’s preferences. The raster cell values intersecting the individual’s current position are extracted for each monthly raster layer (temperature, precipitation, and NDVI). These values are averaged for each environmental variable to represent the local conditions. Next, the averaged environmental values are compared to the individual’s preferences, which are the average of their monthly preference values for each environmental variable. The final yearly movement distance is proportional to the difference between the extracted and preferred values. If the difference is large, the individual moves farther; if the difference is small, the movement is reduced. The computed movement is applied positively or negatively depending on the direction of the difference, enabling individuals to navigate toward areas more closely aligned with their environmental preferences. This method ensures that individuals respond dynamically to environmental gradients while accounting for the species-specific movement limitations.

### Population Modeling

The birth rate within the IBM was simulated based on the life history traits of *T. hudsonicus*. Birth rates were adjusted to reflect variations in reproductive success with age, as reported in previous studies (Descamps et al., 2008). Individuals in the model were assigned as either male or female, with reproduction restricted to females. For a female to successfully reproduce in a given year, a male must be located within approximately 79.8 meters. This distance was calculated as the radius of a circle with an area of 2 ha, which is the average home range size of *T. h. grahamensis*. Reproductive success is highest among females between the ages of 1 and 3 years, with a gradual decline in birth rates as individuals age. *T. hudsonicus* reaches reproductive maturity at 1 year of age (Descamps et al., 2008). To incorporate stochasticity, randomness was introduced to the birth rate calculation. This adjustment ensures variability in reproductive outcomes while constraining the maximum number of individuals capable of giving birth to half the population, reflecting observed population dynamics. This approach enables the model to realistically simulate birth rates across different age classes while incorporating both biological patterns and random variability. The final reproductive model reflects the dynamic and age-dependent nature of *T. hudsonicus* reproduction within a population.

### Mortality and Death Rate

The death rate in the IBM for *T. h. grahamensis* accounts for age-related mortality, infant mortality, maximum age, and population size relative to carrying capacity. This section describes the equations and assumptions used to simulate these dynamics. Randomness was incorporated to better simulate natural variability in mortality. This ensures that mortality probabilities vary stochastically within realistic bounds. Additionally, the final age-adjusted death rate incorporates infant mortality and maximum age adjustments, ensuring biologically realistic mortality patterns. The carrying capacity (*K*) for the *T. h. grahamensis* was estimated at 260 individuals based on population trends observed from 2001 to 2016 (U.S. Fish and Wildlife Service, 2024). This period followed a significant population decline caused by a bark beetle (Scolytinae) infestation between 1998 and 2001 and preceded the Frye Fire of 2017, which led to another marked decline. The final independent death rate (excluding environmental or natural phenomena like wildfire) was calculated by multiplying the age-adjusted death rate (*C*_*d*_) by the proximity factor (*F*_*c*_): Final Death Rate = *C*_*d*_ × *F*_*p*_. Randomness was further incorporated into the death rate to simulate real-world variability, except for individuals at the maximum age (*a* = 10), where the death probability was fixed at 100%. If the population size reaches 5% of the original population, the death probability doubles to simulate the increased vulnerability of small populations. This approach ensures a realistic simulation of mortality patterns, accounting for biological constraints and environmental pressures specific to *T. h. grahamensis*.

#### Mortality Modeling Due to Climatic Factors

Environmental factors affecting mortality are calculated by comparing the individual’s location to the species’ preferred environmental ranges. The following equations quantify the likelihood of mortality due to precipitation, temperature, and NDVI values:

The output of these equations is weighted to reflect the relative importance of each factor for *T. h. grahamensis* survival. Precipitation and NDVI, which influence food and shelter availability, are given greater weights than temperature. The final environmental factor (*F*_*E*_) determining mortality due to climatic conditions is calculated as:

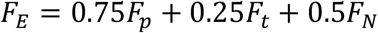

This approach ensures that mortality risk is driven primarily by the most critical environmental variables—precipitation and NDVI—while incorporating the influence of temperature. By weighting these factors and incorporating species-specific preferences, the model accurately simulates the impact of unfavorable climatic conditions on the mortality of the *T. h. grahamensis*.

#### Environmental Disturbance Simulation in the Model

Wildfires have become an increasingly significant threat due to climate change and historical wildfire suppression policies (Halofsky, 2020). To account for the impacts of wildfires, diseases, and pest infestations with the *T. h. grahamensis* population, large-scale disturbances were incorporated into the simulation based on environmental disturbance history in the Pinaleño Mountains (*see Environmental Disturbances in Supplemental Information*). The wildfire simulation occurred using the following constraints: fires ranged from 405 ha to a maximum of 16,187 ha (determined using a uniform distribution) and the fire’s impact was modeled as a polygon originating from a randomly selected central point. Insect Infestations were modeled to encapsulate the entire high-elevation vegetation region (~28,328 ha), but with a lower death rate than wildfires based on infestation history in the Pinaleño Mountains. Based on recent trends in the Pinaleño Mountains, the model assigns a 10% annual probability of triggering a wildfire event with areas with higher NDVI values (indicating denser vegetation) having a greater likelihood of triggering wildfires due to the increased availability of fuel for burning.

Individuals located within the wildfire radius have a 70% probability of mortality. This probability reflects the potential for direct death due to the fire as well as indirect impacts, such as loss of shelter and food resources. The probability of survival also accounts for variability in fire intensity; some fires leave trees and vegetation partially intact, reducing their impact on critical resources (Snow, 2022). Consequently, the chance of death is not set to 100%. By incorporating wildfire dynamics, the model reflects the role of natural disturbances in shaping population survival and distribution.

#### Climate Change Implementation in the Individual-Based Model

The IBM incorporates climate change projections for temperature, precipitation, and NDVI to simulate potential impacts on the *T. h. grahamensis* population. These changes are derived from authoritative sources, such as the National Climate Assessment (2014), and previous studies. Projections account for gradual changes over time and include worst-case scenarios to assess the implications for the Madrean Archipelago’s sky island ecosystems. Temperature projections are based on the National Climate Assessment (2014), which predicts a rise of 2.5 to 5.5°F (1.4 to 3.0°C) in the southwestern United States by 2041–2070 under maximum emissions scenarios. See *Climate Change* section in the Supplemental Materials for specific yearly change calculations. Precipitation projections are also sourced from the National Climate Assessment (2014), which predicts a 5–20% decrease in precipitation by 2100 under maximum emissions scenarios. Changes in NDVI are based on prior studies conducted on sky islands, where lower-elevation habitats exhibited an average NDVI decline of 13.17% over 11 years. The studies were completed in May 2024 using ArcGIS Pro to analyze changes in NDVI over an 11-year period.

Annual assessments focused on various habitats and elevations, leveraging Landsat raster data to evaluate shifts in average NDVI across these regions. The analysis revealed that higher elevations with greater NDVI values experienced minimal change, whereas lower elevations with lower NDVI values exhibited the most pronounced declines in NDVI (**Supplemental Materials)**. To reflect the differential impacts of climate change across elevations, the model applies larger NDVI reductions in areas with lower initial NDVI values and smaller reductions in areas with higher initial NDVI values.

##### Simulation Scenarios

The IBM implements these climate and environmental changes alongside simulations with no climate change to serve as a baseline. By comparing these scenarios, the model assesses worst- and-best-case outcomes for the *T. h. grahamensis* population and other species inhabiting the Madrean Archipelago’s sky islands. This approach provides insights into the potential ecological consequences of climate change in these unique and vulnerable ecosystems. The Python programming language was used within Jupyter Notebooks for the simulation and the code is available on GitHub (see **References** section)

## Results and Discussion

### Model Assessment

Identifying the appropriate habitat boundaries was essential for creating agent-based models. We found that the model effectively delineates sky island boundaries (**p < 0.01, Figure 2**).

**Figure 2.**
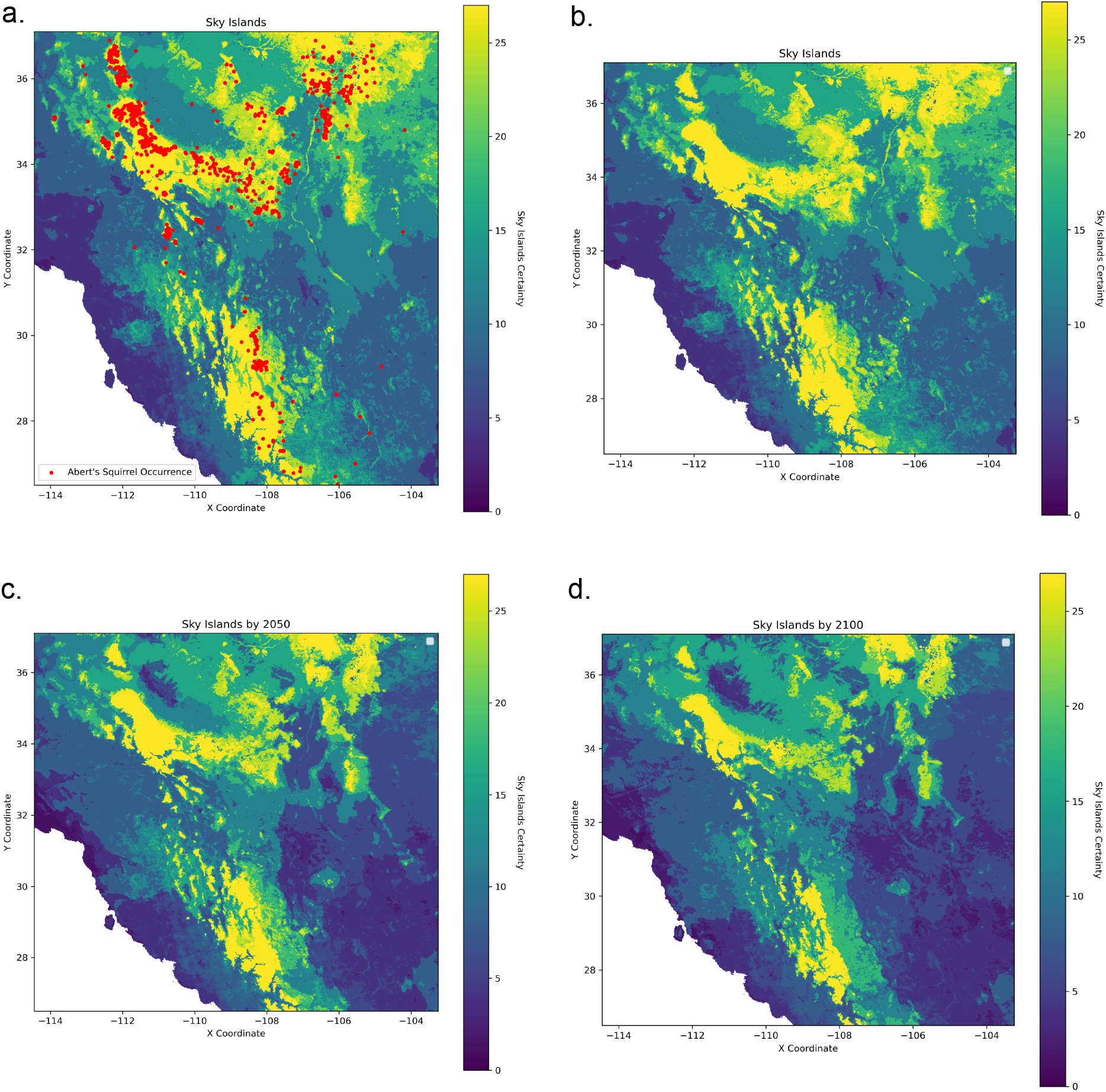
Sky Island habitat suitability (A) Occurance points for *T. h. grahamensis* (B) Current Sky Islant Habitat (C) Sky Island Habitat in 2050 (D) Sky Island Habitat in 2100.

**Figure 3.**
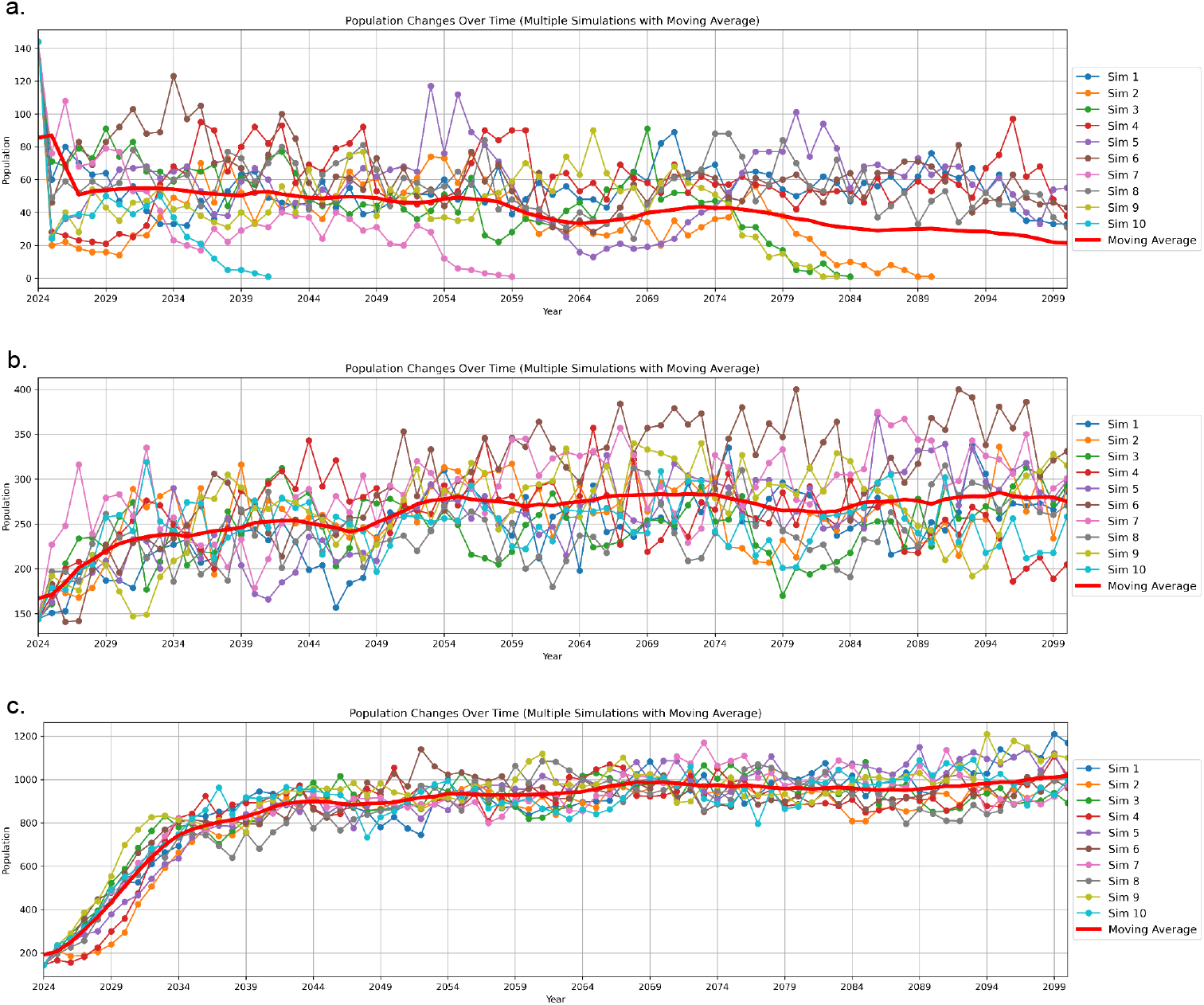
Sensitivity Analysis of Proximity Exponents in Population Dynamics. This figure represents a sensitivity analysis conducted to determine the optimal proximity exponent for simulations incorporating climate change and environmental disturbances. However, these disturbances are excluded from the current analysis. *T. h. grahamensis* has an estimated carrying capacity of approximately 260 individuals within the Pinaleño Range under conditions with minimal climate change and limited environmental disturbances. The proximity exponent is a user-defined parameter that adjusts the probability of mortality exponentially in relation to population size. Mortality probability is inversely proportional to the average distance between individuals, meaning closer proximity leads to higher mortality rates (see Supplemental Materials for details). a) This panel shows 10 simulations with a proximity exponent of −6.5. In this scenario, the probability of death reaches 100% at a population of 147 individuals if distance is disregarded. The resulting carrying capacity is approximately 40 individuals, far below the target value of 260, invalidating this exponent for final scenario simulations. b) This panel illustrates 10 simulations using a proximity exponent of −8. The carrying capacity aligns closely with the target of 260 individuals, consistent with observed population data under minimal climate change and environmental disturbance. As a result, this exponent was selected for use in the final scenario simulations. c) This panel depicts 10 simulations with a proximity exponent of −9.5. Here, the probability of death reaches 100% at a population of 1,468 individuals. The resulting carrying capacity of approximately 1,000 individuals significantly exceeds the target of 260, eliminating this exponent from consideration. Following 30 simulations testing three proximity exponents, −8 was identified as the most accurate in replicating observed population dynamics. At this value, the probability of death, independent of distance, is 18.6%, yielding a carrying capacity consistent with empirical data under conditions without environmental disturbances. This sensitivity analysis informed the selection of −8 for final scenario simulations.

**Figure 4.**
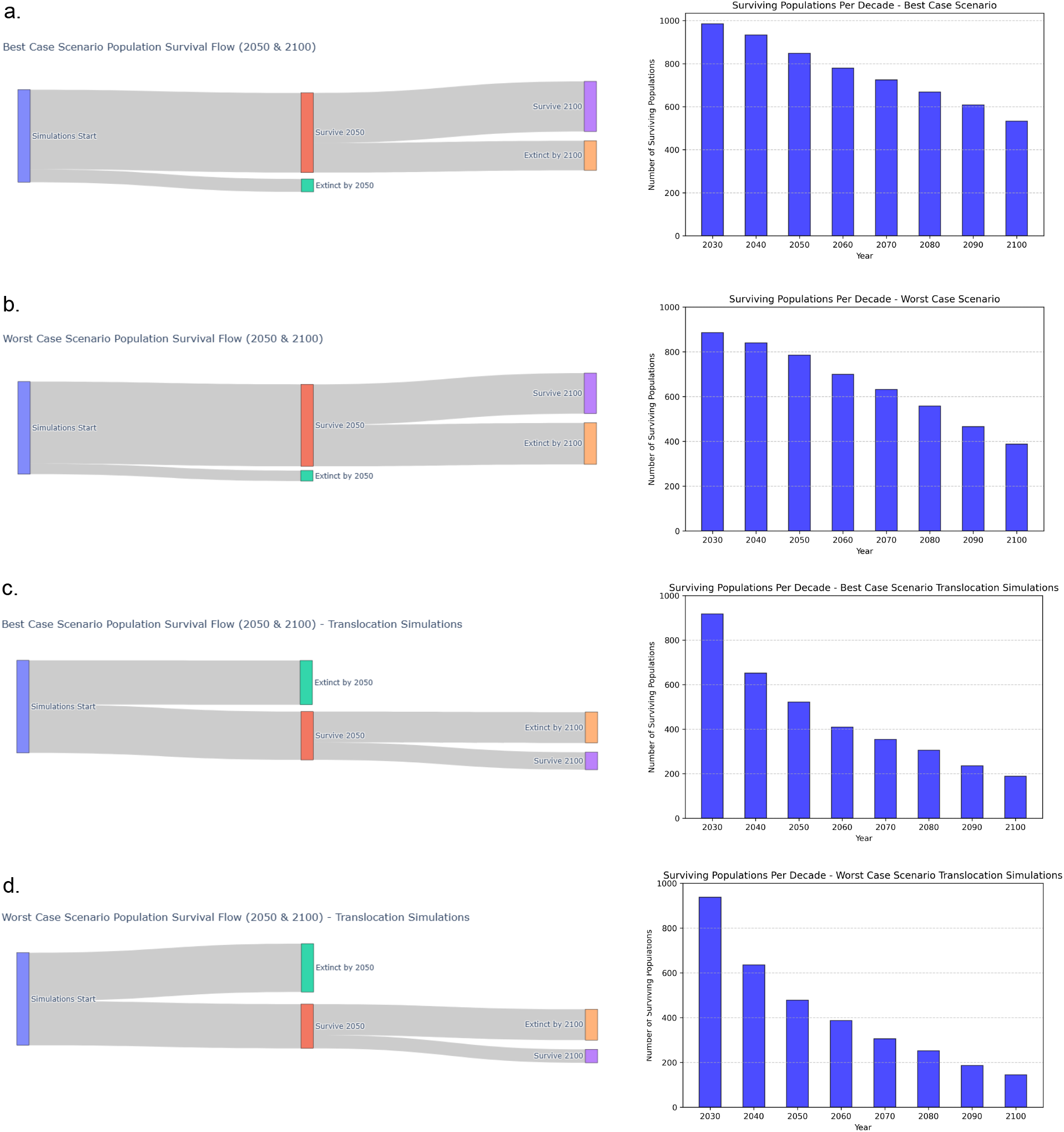
Final Scenario Simulations Incorporating Climate Change and Environmental Disturbances. The final scenario simulations integrate both best- and worst-case climate change models, along with the most significant environmental disturbances affecting the Pinaleño Range, specifically wildfires and insect infestations. A total of 1,000 simulations were conducted for each climate model to compute an accurate moving average of T. h. grahamensis populations through the year 2100. Beyond estimating population trends, these simulations were analyzed for potential extinction events and challenges in population recovery. Results indicate that frequent and successive environmental disturbances contribute to significant population declines, thereby increasing the probability of extinction. Additionally, these disturbances may displace individuals, forcing populations to inhabit suboptimal environmental conditions, which further exacerbates survival challenges. (A) Best-Case Climate Scenario Model Simulations Of the 1,000 simulations conducted under the best-case climate scenario, 541 populations persisted beyond 2100, suggesting a 46% probability of extinction by the end of the century. Additionally, 139 populations were lost before 2050. Rapid population increases following environmental disturbances, particularly wildfires, played a crucial role in mitigating early extinctions. (B) Worst-Case Climate Scenario Model Simulations In contrast, the worst-case climate scenario resulted in a substantial number of population extinctions before 2100. Only 437 simulations contained populations that survived until 2100, indicating a mere 44% probability of persistence by the end of the century. Furthermore, only 886 populations survived until 2050. These results suggest that under the worst-case scenario, T. h. grahamensis faces a greater than 50% likelihood of extinction before 2100, with major environmental disturbances being the primary driver of population decline. (C) Best-Case Translocation Model Simulations Under the best-case climate scenario, translocation of 50 individuals to the Chiricahua Mountains in southeastern Arizona yielded poor survival outcomes. By 2050, only 478 simulations contained surviving populations, with survival dropping to just 145 by 2100. These findings indicate that even under optimal climatic conditions, translocation offers only a minimal chance of long-term survival for this endangered species, suggesting that it may not be a viable conservation strategy for T. h. grahamensis. (D) Worst-Case Translocation Model Simulations Under the worst-case climate scenario, translocation efforts resulted in even higher extinction probabilities. By 2100, only 145 simulations contained surviving populations, corresponding to an 89% probability of extinction before the end of the century. Additionally, only 478 simulations retained populations until 2050. The outcomes depicted in Graphs c. and d. likely stem from the lower elevations and consequently higher overall temperatures of the Chiricahua Range. The Pinaleño Mountains, home to the highest peak in the Madrean Archipelago (Mount Graham, 3,267 meters or 10,720 feet), are situated further north than larger sky islands such as the Chiricahua and Huachuca ranges. Translocation to areas along the Mogollon Rim may benefit overall red squirrel populations but would compromise the evolutionary uniqueness of T. h. grahamensis.

Additionally, the results align with known ecological distributions of squirrel populations, further supporting the model’s validity.

#### Reconstruction of Historical Populations

##### Climate change simulations

These simulations include the projected worst- and best-case scenario of changes from 2024 to 2100 regarding temperature, precipitation, and NDVI according to the 2014 National Climate Assessment and previous studies (see *Supplemental Information*). Based on these future projections, the environmental factors involved within this simulation changed over time. Similar to what is currently observed in field studies, *T. h. grahamensis* populations are forced to higher elevations where temperatures, precipitation, and vegetation will be less affected (Rushton et al., 2018).

Extinction within *T. h. grahamensis* is primarily driven by significant environmental disturbances affecting the population. Historical disturbances, such as the bark beetle infestation in 1998 and the Frye Fire in 2017, caused substantial population declines of approximately 55% and 86%, respectively. While *T. h. grahamensis* populations have previously demonstrated the ability to recover following such disturbances, climate change model simulations indicate that this resilience has diminished. This reduced capacity for recovery is likely due to displacement of individuals into less suitable habitats, increasing mortality rates and reducing opportunities for population growth. Additionally, these environmental disturbances exert prolonged generational effects due to the destruction of vegetation, which diminishes both shelter and food resources.

Although insect infestations, such as those caused by bark beetles, do not directly result in significant mortality, their ecological impact persists for decades, as forest recovery following such events is markedly slow. In contrast, wildfires often result in immediate and severe environmental impacts and high mortality rates; however, they may also promote rapid forest regrowth through the fertilizing effects of ash and other burned materials (Santín, 2016). This regrowth can mitigate the duration of wildfire-induced disturbances, facilitating population recovery, as evidenced in the final scenario simulation graphs. Simulations show that populations typically recover relatively quickly following wildfires, despite the initial drastic declines in numbers. Complete extinction due to wildfires was only observed in scenarios where fires occurred in rapid succession—an improbable scenario for this region. Importantly, climate change models suggest that population shifts are not drastically influenced by gradual changes in environmental conditions, particularly under best-case scenarios. Instead, the challenges to population viability are predominantly associated with increasingly frequent and destructive events driven by changing environmental conditions. Warmer and drier climates exacerbate the prevalence and severity of wildfires and promote overpopulation of harmful species, such as bark beetles, which thrive in higher temperatures.

### Species Protection Via Translocation

Translocation is the process by which a small population of a species is relocated to a new area to prevent the species from reaching extinction. In upcoming years, according to the population simulations, it is likely some sort of conservation initiative will need to be launched, and this could be a possibility. Assisted migration has historically proven to be an effective conservation strategy, and its implementation could significantly benefit species within the Madrean Archipelago (Camacho, 2010). It is possible to introduce a population of *T. h. grahamensis* to a larger forested area, like the Mogollon Rim region of Arizona. Unfortunately, these areas include a much greater diversity of species, leading to more competition, and may inhibit the genetic uniqueness of *T. h. grahamensis*. A possibility for supporting the population for the foreseeable future, while also protecting their evolutionary developments could be to move a small population of the species to a larger sky-island.

The Chiricahua Range is a mountain range located approximately 100 kilometers south-southeast of the Pinaleño Range. The Chiricahua Mountains cover approximately 1.5 times the area of the Pinaleño Mountains, spanning roughly 1,172 square kilometers (AZIBA, 2009), while the Pinaleño Range encompasses only 777 square kilometers (Curiel, 2009). This could make the region a possibility for population reconstruction within the next century. A sensitivity analysis found minimum number of *T. h. grahamensis* individuals which could be relocated to the Chiricahua Mountains and exhibit population growth. The minimum population to be relocated was ~50 individuals, which were then all placed at the same location to simulate conservationists releasing the individuals. After performing another sensitivity analysis for selection of a release location, the Herb Martyr Campground was selected for its abundance of large pine tree species and higher elevation, exhibiting environmental conditions similar to those *T. h. grahamensis* prefer.

After conducting 1,000 simulations under both best- and worst-case climate scenarios, our results indicate that translocation may not be a viable conservation strategy for the species. Alarmingly, 575 out of 1,000 simulated populations experienced extinction by 2100, suggesting a significantly higher probability of extinction compared to the naturally occurring population in the Pinaleño Mountains. *T. h. grahamensis* already maintains a relatively small population size compared to many other small mammal species, which limits the number of individuals available for translocation and further exacerbates the risk of extinction. Moreover, although the Chiricahua Range is larger and contains a greater extent of forested habitat, it has a lower average elevation than the Pinaleño Range, resulting in slightly reduced precipitation and higher temperatures. Additionally, the frequency of wildfires and insect infestations in the Chiricahua Range is comparable to that of the Pinaleño Range, offering no substantial relief from these major ecological stressors. If translocation were done to areas where there would be no interspecific hybridization, the species may remain as it is, however if it is translocated to areas where there is substantial hybridization new species may emerge.

## Conclusion

According to the best-case scenario climate model, conservation efforts are essential to support the *T. h. grahamensis* population during this century. However, it is unlikely that the entire population will face extinction within the next 20 years. The species demonstrates adaptability to severe environmental changes, as evidenced by population recovery following drastic climate events. Nonetheless, by 2100, there remains an approximately 25% likelihood of extinction. Climate change drives vegetation loss at lower elevations, with these effects progressively shifting to higher elevations as the climate becomes warmer and drier. This results in reduced vegetation for shelter and food, forcing the population into higher elevations and smaller habitats. Consequently, competition increases among individuals and with other species, such as *Sciurus aberti*, as resources become scarcer. This intensified competition explains the population’s rapid recovery after environmental disturbances: die-offs reduce competition, allowing for unimpeded population growth once the habitat naturally recovers, particularly following wildfires. The conservation of *T. h. grahamensis* is of paramount importance due to its small population size and restricted endemic range. However, population simulations and historical records (Koprowski, 2008) suggest that the species is more resilient than previously believed. Its capacity to endure relatively arid and warm conditions, coupled with rapid recovery rates, simplifies conservation efforts. A key strategy for preserving the species involves reducing the occurrence of large crown fires, such as the 2017 Frye Fire, as consecutive large-scale disturbances are the most common drivers of extinction. Implementing prescribed burns in the Pinaleño Mountains could mitigate this risk while promoting vegetation regrowth through ash fertilization, supporting both the species and its habitat.

## Supplemental Materials ODD description

This supplement can be used as a template for writing ODD model descriptions. It contains Section 3 of the manuscript. After reading the explanations and typing the answers to the question, ODD users should have a clear and complete ODD model description of their individual- or agent-based models. Questions and explanations should, of course, be deleted then.

### Overview, Design concepts, Details Protocol for the Sky Island Species Model

#### 1. Purpose

- **Question:** What is the purpose of the model?
  - **Answer:** The model aims to predict the population dynamics (growth, decline, movement) of the Mount Graham Red Squirrel (*Tamiasciurus hudsonicus grahamensis*) under current and future climate scenarios. The goal is to determine if, when, and how this species might face extinction due to environmental changes, providing actionable insights for conservation planning.
  - **Explanation:** This model addresses ecological questions concerning the survival of isolated endangered species in sky island ecosystems under climate change pressures. It helps bridge theoretical ecological principles with applied conservation strategies.

#### 2. Entities, State Variables, and Scales

- **Entities:**
  - Individual squirrels: characterized by attributes such as ID, age, sex, location, fitness, and adaptive traits.
  - Grid cells (representing habitat): characterized by elevation, vegetation type, NDVI, temperature, precipitation, and resource availability.
  - Environment: represents global factors such as overall climate trends and stochastic events like droughts.
- **State Variables:**
  - Squirrels: age, reproductive status, location, energy reserves, and survival probability.
  - Grid cells: vegetation cover, resource density, and suitability index.
  - Environment: temperature and precipitation time series.
- **Scales:**
  - Temporal: Each time step represents one year; simulations run from 2024 to 2100.
  - Spatial: Grid cells represent 1 km^2^, covering the sky island ecosystems in the Pinaleño Range, Arizona, USA.

#### 3. Process Overview and Scheduling

- **Who does what:**
  1. Squirrels move, forage, reproduce, and die based on individual traits and environmental conditions.
  2. The environment updates habitat suitability indices based on climatic inputs and vegetation dynamics.
- **Order of operations:**
  1. Update environmental conditions (temperature, precipitation).
  2. Calculate habitat suitability for each grid cell.
  3. Simulate squirrel movement and foraging.
  4. Simulate reproduction for eligible individuals.
  5. Update population demographics (age, mortality).
- **Time:** Modeled in discrete yearly steps.

#### 4. Design Concepts

- **Basic Principles:** Based on ecological theories of species distribution, resource competition, and climate adaptation. The model further incorporates climate resilience strategies to evaluate the long-term survival of the species.
- **Emergence:** Population trends and extinction events emerge from individual behaviors and environmental interactions. These patterns provide insights into critical thresholds and tipping points in ecosystem dynamics.
- **Adaptation:** Squirrels adapt their movement and foraging strategies to maximize resource intake and survival. These strategies are constrained by environmental heterogeneity and agent competition.
- **Objectives:** Individuals aim to survive, reproduce, and maximize fitness within their environmental constraints. This ensures that individual-level decisions drive population-level patterns.
- **Learning:** Not explicitly modeled; behaviors are predefined based on ecological rules. However, potential pathways for incorporating learning mechanisms are identified for future model iterations.
- **Prediction:** Implicit in the decision-making processes, e.g., selecting habitats based on resource availability. These predictions are based on resource gradients and environmental stability.
- **Sensing:** Individuals sense local environmental conditions and resource availability. Sensing mechanisms allow for real-time adjustments to movement and foraging behaviors.
- **Interaction:** Competition for limited resources within grid cells. Interactions also influence spatial distribution patterns and territoriality.
- **Stochasticity:** Introduced in processes like mortality, reproduction, and climatic variations. This randomness reflects real-world variability and enhances model robustness.
- **Collectives:** Not modeled explicitly but implied in habitat-level dynamics. Potential collective behaviors like group foraging could be explored in future model extensions.
- **Observation:** Outputs include population size, extinction probabilities, and habitat use over time. These metrics are validated against empirical data to assess model accuracy.

#### 5. Initialization

- **Initial State:**
  - Population: 144 individuals distributed within the suitable habitat of Mount Graham.
  - Environment: Habitat suitability maps generated from historical climate data.
  - Grid cells: Initialized with vegetation, temperature, and precipitation values from remote sensing and historical datasets.
- **Variability:** Initialization parameters can vary between simulation runs to test different scenarios.

#### 6. Input Data

- **Data Sources:**
  - Climate: PRISM Climate Group dataset for historical temperature and precipitation; CMIP6 projections for future climate scenarios.
  - Vegetation: MODIS NDVI (Normalized Difference Vegetation Index) data from NASA Earth Observing System.
  - Species distribution: Occurrence data from GBIF (Global Biodiversity Information Facility) for Mount Graham Red Squirrel.
  - Elevation: SRTM (Shuttle Radar Topography Mission) elevation data for generating terrain features.
  - Land cover: USGS National Land Cover Database (NLCD) for vegetation type and habitat classification.
- **Representation:** Inputs drive habitat suitability calculations and influence individual behaviors and population dynamics.

#### 7. Submodels

- **Habitat Suitability:**
  - Combines temperature, precipitation, and NDVI data to calculate habitat suitability indices.
- **Foraging and Movement:**
  - Based on energy optimization and resource availability within grid cells.
- **Reproduction:**
  - Probability based on age, energy reserves, and habitat quality.
- **Mortality:**
  - Stochastic function of age, energy reserves, and environmental stressors.
- **Climate Projections:**
  - Incorporates external climate models to simulate future conditions. See Table 1 in main text for full details.

#### Equations

##### Table 1 - Age-Based Birth Probability

Where:

- *C*_*b*_: Probability of giving birth
- *a*: Individual’s current age
- *a*_*m*_: Maximum age an individual can reach
- *e*: Euler’s number

##### Table 1 - Age-Based Mortality Probability

Where:

- *C*_*d*_: Probability of death
- *a*: Individual’s current age
- *a*_*l*_: Species’ average lifespan (5 years)
- *e*: Euler’s number

##### Table 1 - Population-Based Mortality Rate

Where:

- *exp*_*p*_ = user-defined proximity exponent
- *P* = current population
- *w*_*p*_ = proximity weight

#### Stand-alone equations

Comparison of Environmental Conditions to Individual Preferences

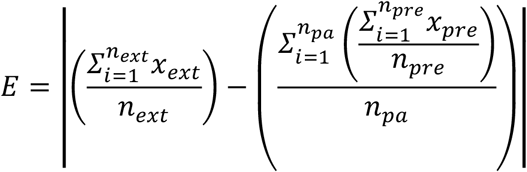

Where:

- *E*: Difference between the extracted value and the preferred value
- *x*_*ext*_: Extracted environmental value
- *n*_*ext*_: Number of extracted values
- *x*_*pre*_: Individual’s preference value
- *n*_*pre*_: Number of preference values
- *n*_*pa*_: Number of averaged preference values

##### Yearly Movement

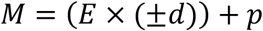

Where:

- *M*: Total yearly movement
- *E*: Difference between the extracted and preferred values
- *d*: Maximum yearly movement distance (679.8 meters)
- *p*: Current position

##### Proximity Factor

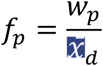

Where:

- *w*_*p*_ = proximity weight
- 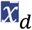 = average distance between individuals
- *f*_*p*_ = proximity factor

##### Environmental Preferences

#### Precipitation Factor (*F*_*p*_)

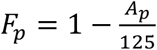, with a minimum value of 0.

- *A*_*p*_: Average precipitation preference of the individual

#### Temperature Factor (*F*_*t*_)

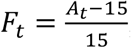, with a maximum value of 1.

- *A*_*t*_: Average temperature preference of the individual

#### NDVI Factor (*F*_*N*_)

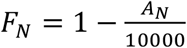, with a minimum value of 0.

- *A*_*N*_: Average NDVI preference of the individual

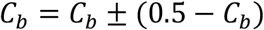

#### Geospatial Information

The WGS84 (EPSG:4326) coordinate system was used to simulate the Individual-Based Model. Distances were converted to decimal degrees, adjusted for latitude. A latitude of **32.7°** was used as the conversion factor, reflecting the central location of the study area.

#### Environmental Disturbances

##### Historical Wildfire Trends in the Pinaleño Mountains

The Pinaleño Mountains have experienced several large-scale wildfires in recent decades, as recorded by the University of Arizona. Notable events include:

- **Clark Peak Fire (1996):** Burned approximately 6,300 acres.
- **Nutfall Complex Fire (2004):** Burned approximately 29,200 acres.
- **Frye Fire (2017):** Burned approximately 48,443 acres.

Wildfires are also becoming larger and more frequent in broader contexts. For instance, in California, **18 of the 20 largest wildfires** between 1980 and 2021 occurred between 2003 and 2021, with **half of these occurring in 2020 and 2021 alone** (MacDonald, 2003). These trends indicate that large-scale wildfires are becoming a persistent challenge.

#### Climate change

##### Temperature

- The midpoint temperature increase is calculated as **4°F (2.2°C)**.
- The average annual temperature increase is derived as:

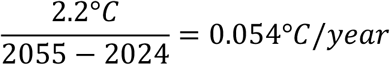

##### Precipitation

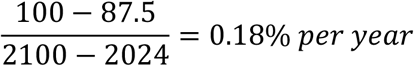

This annual reduction (0.18%) is applied uniformly across the study area, simulating a gradual decline in water availability over time.

##### NDVI

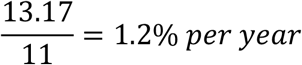

This equation only applies to areas of lower NDVI, reflecting areas of lower elevation, which tend to be targeted first in desertification events.

**Figure S1.**
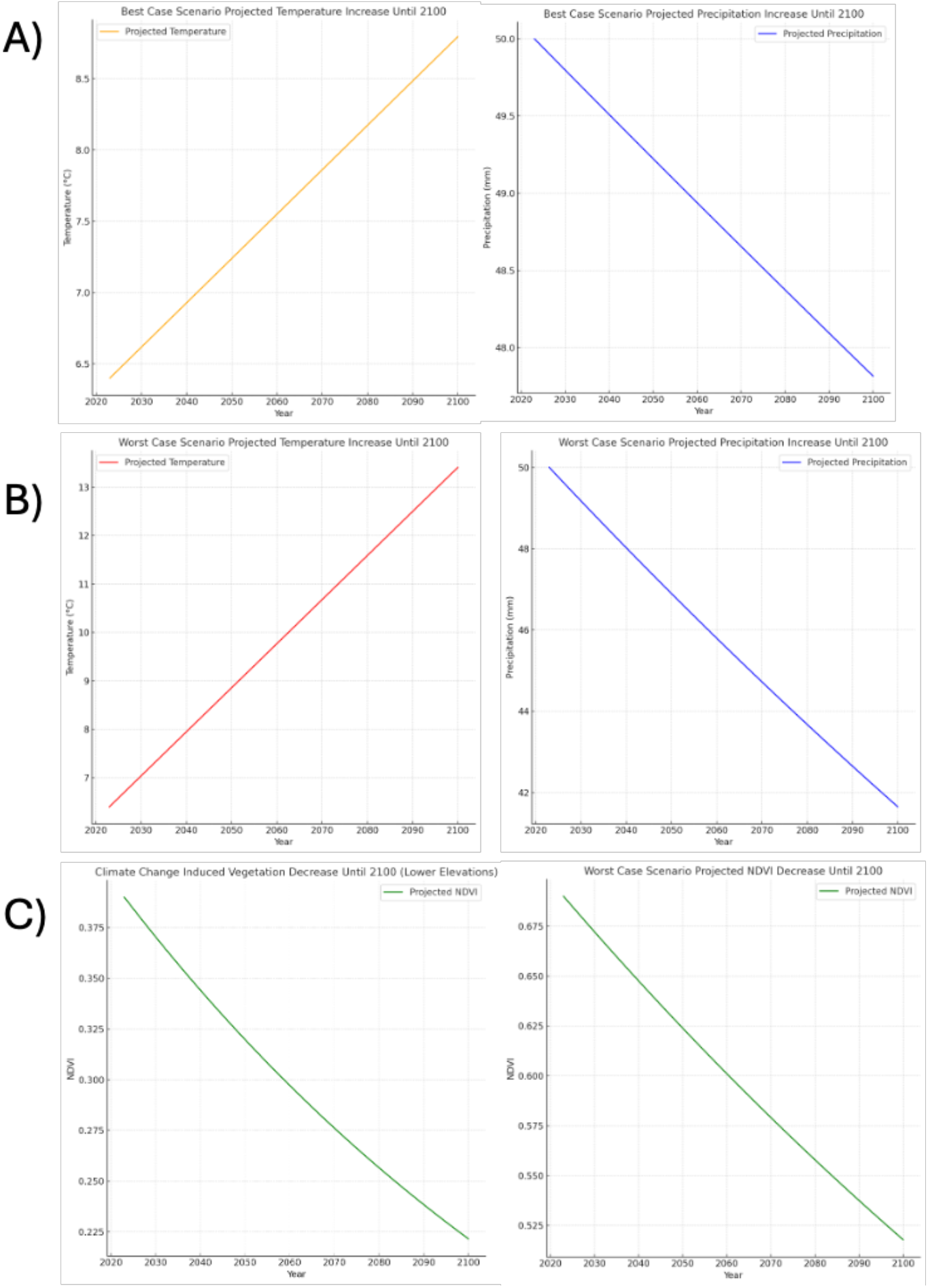
This figure illustrates the projected changes in temperature, precipitation, and NDVI (Normalized Difference Vegetation Index) under future climate scenarios based on IPCC predictions and previous studies of sky islands within the Madrean Archipelago. These temporal changes are depicted for both best- and worst-case scenarios, representing the potential range of environmental impacts. Final scenario simulations will utilize these projections to estimate the minimum and maximum population changes for *T. h. grahamensis* by 2100. (**a)** The first panel displays temperature, precipitation, and NDVI projections for a best-case scenario according to IPCC climate predictions. Under this scenario, temperature is projected to increase by 1.9°C and precipitation to decrease by 4.5% by 2100. Simulations incorporating these parameters represent the least impactful climate conditions, effectively modeling the minimal population effects possible by 2100.**(b)** The second panel presents projections for a worst-case scenario, where temperature is expected to increase by 6.9°C and precipitation to decrease by 18% by 2100. These parameters characterize the most severe climate impacts, modeling the maximum potential population declines by 2100. **(c)** Changes in NDVI are consistent across both scenarios, as they are derived from recent vegetation data and do not include direct predictions of climate-driven changes in NDVI for this region. Additionally, NDVI adjustments vary across elevations: lower elevations, with initially lower NDVI values, experience more pronounced declines due to desertification, with these effects progressively shifting to higher elevations over time.Temperature and precipitation graphs in both panels are initialized with average annual values preferred by *T. h. grahamensis* based on monthly environmental preferences. The initial temperature value is 6.4°C, while the initial precipitation value is 50 millimeters. NDVI graphs include separate trajectories for low- and high-elevation values to reflect spatial variation. The lower NDVI value graph begins at the 25th percentile of NDVI preferences (0.39), while the higher NDVI value graph begins at the 75th percentile (0.69). Importantly, all changes are applied to monthly temperature, precipitation, and NDVI rasters, ensuring dynamic updates throughout the simulation period.

## Notes

### Competing Interest Statement

The authors have declared no competing interest.

